# Comparing the limbic-frontal connectome across the primate order: conservation of connections and implications for translational neuroscience

**DOI:** 10.1101/2024.03.06.583735

**Authors:** Davide Folloni, Lea Roumazeilles, Katherine L Bryant, Paul R Manger, Mads F Bertelsen, Alexandre A Khrapitchev, Peter H Rudebeck, Rogier B Mars

## Abstract

The interaction of the limbic system and frontal cortex of the primate brain is important in many affective behaviors. For this reason, it is heavily implicated in a number of psychiatric conditions. This system is often studied in the macaque monkey, the most largely-used non-human primate model species. However, how evolutionary conserved this system is and how well results obtained in any model species translate to the human can only be understood by studying its organization across the primate order. Here, we present an investigation of the topology of limbic-frontal connections across seven species, representing all major branches of the primate family tree. We show that dichotomous organization of amydalofugal and uncinate connections with frontal cortex is conserved across all species. Subgenual connectivity of the cingulum bundle, however, seems less prominent in prosimian and New World monkey brains. These results inform both translational neuroscience and primate brain evolution.

## INTRODUCTION

The limbic system and frontal cortex form an interconnected system of brain areas that are affected in disorders of mood, motivation, and anxiety ^1–3^. Reciprocal connectivity between the parts that form this system are essential to support their roles in associative learning, decision-making and behavioral control ^4–6^. As such, these connections are often the target of clinical interventions such as deep brain stimulation ^7–9^. Advances in non-invasive stimulation of deep cortical structures ^10–13^ mean that functional interactions between connectional systems can potentially be targeted without surgical interventions. However, to advantageously and judiciously use these advances, a better understanding of the limbic-frontal connectome is required.

Much of our knowledge of these systems is based on invasive work in non-human animals, using techniques that cannot be ethically/legally applied to humans. The development of neuroimaging techniques that can be applied to both humans and non-human animals with increasing levels of resolution has enabled formal tests of how well results obtained in model species translate to humans ^14^. For example, comparative diffusion MRI-based tractography work has concentrated on assessing the principles of frontal cortex organization and connectivity in humans and macaque monkeys ^15–17^. Recent investigations demonstrate how such methods can also be used to identify even very specific limbic-frontal connections. For instance, the amygdalofugal pathway, a small pathway running between the amygdala and frontal cortex (PFC) ^18–21^, could be detected in humans using tractography, as long as the approach is guided by anatomical knowledge obtained in macaques using both invasive tracers and tractography ^22^. Comparative neuroimaging can thus be a valuable tool for investigating how limbic-frontal circuits vary across species.

Neuroimaging has the additional advantage that it can be applied to a larger number of species than can often be used in comparative neuroanatomical studies. Comparisons do not have to be limited to studies of humans and macaques and can be extended to a broader range of primate species, allowing the characterization of patterns of evolutionary consistency or change. Understanding such patterns will lead to a better understanding of how well results obtained in any models species are likely to generalize to the species of interest, which in most cases is the human. The increased availability of MRI datasets from non-human primates ^23,24^, combined with new analytical frameworks ^25^, make such studies feasible.

Frontal cortex is thought to be quite variable across primate species, both in term of its relative size and connectivity with other brain regions ^26–28^. Therefore, we set out to compare the major pathways that connect to frontal cortex from the temporal lobes and other parts of the brain. It is possible that organizational variances of these frontal cortical areas among primates may be associated with differently organized patterns of connectivity. Alternatively, the course and organization of the amygdala-cingulate-prefrontal circuits may be conserved across primate species. To test these possibilities, we reconstructed the cingulum bundle (CB), ventral amygdalofugal pathway (AmF), and the uncinate fascicle (UF) across an unprecedented range of primate species using probabilistic tractography (**Figure 1**). We observed that, at the level of diffusion-tractography estimates of anatomical connectivity, the amygdala-cingulate-prefrontal circuits are similar in their course and overall pattern of organization not only in anthropoid brains, but also in prosimians.

**Figure 1.**
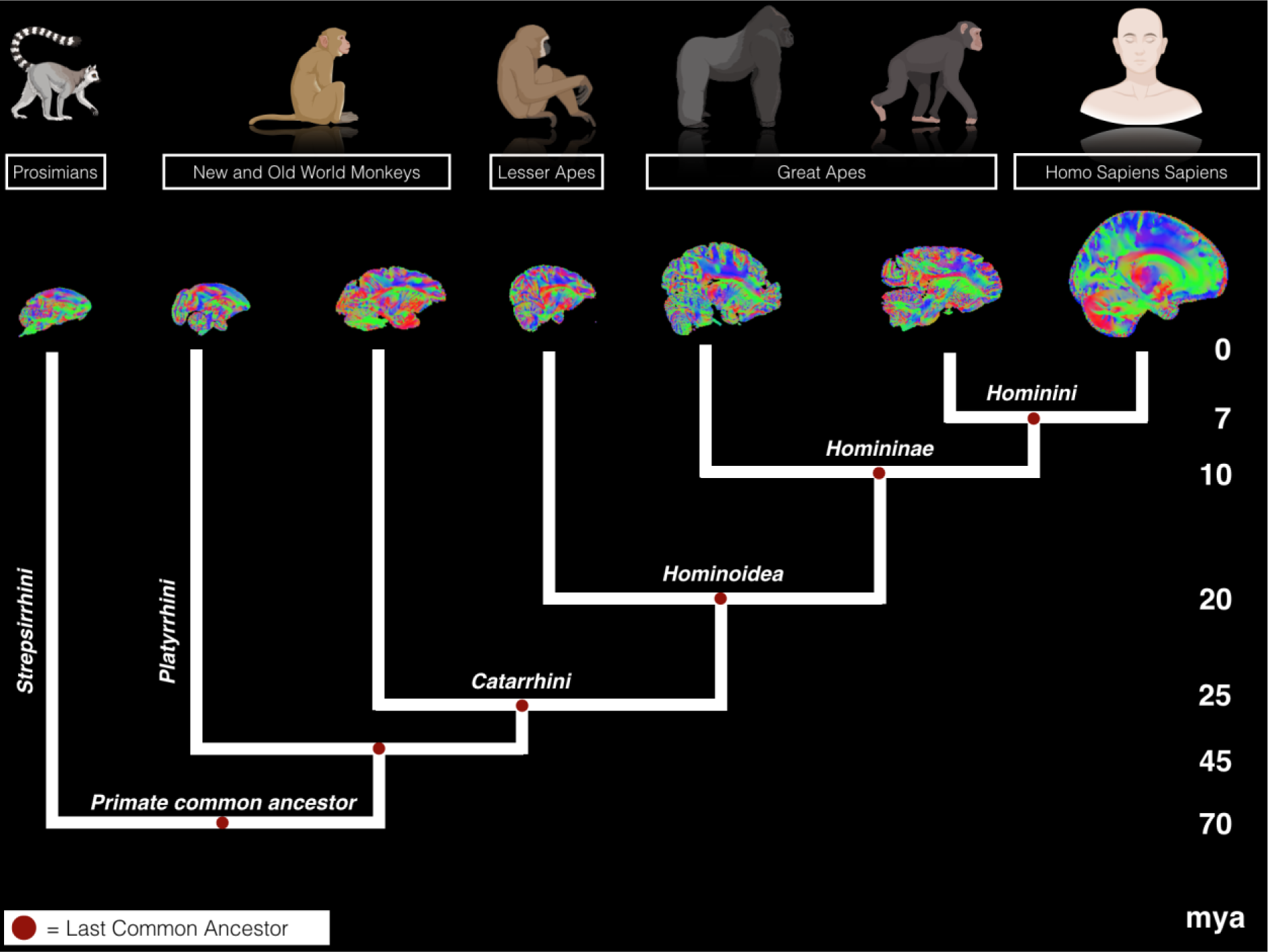
Primate family tree. Phylogram showing evolutionary relationships of prosimians, Old and New World monkeys, lesser and great apes, and humans ^58,60–62^. Brains are not to scale. Mya = million years ago.

## METHODS

### Data acquisition and preprocessing

#### Human data

Human *in vivo* diffusion MRI data were provided by the Human Connectome Project (HCP), WU-Minn Consortium (Principal Investigators: David Van Essen and Kamil Ugurbil; 1U54MH091657) funded by the 16 NIH Institutes and Centers that support the NIH Blueprint for Neuroscience Research; and by the McDonnell Center for Systems Neuroscience at Washington University ^29^. The minimally preprocessed datasets of the first twenty-four subjects from the Q2 public data release were used (age range 22-35 years; thirteen females). Data acquisition and preprocessing methods are detailed in Ugurbil et al. ^30^, Sotiropoulos et al. ^31^, and Glasser et al. ^32^. The diffusion data were collected at a 1.25 mm isotropic resolution across the entire brain on a customized 3T Siemens Skyra scanner using a monopolar Stejskal-Tanner diffusion scheme with a slice-accelerated EPI readout. Sampling in q-space included 3 shells at b = 1000, 2000, and 3000 s/mm^2^. For each shell 90 diffusion gradient directions and 6 non-diffusion weighted images (b = 0 s/mm^2^) were acquired with reversed phase-encoding direction for TOPUP distortion correction ^33^. Voxel-wise model fitting of diffusion orientations was performed using FSL’s bedpostX to fit a crossing fiber model to the data ^34^. A multi-shell extension was used to reduce overfitting of crossing fibers due to non-monoexponential diffusion decay ^35^. Up to three fiber orientations per voxel were allowed.

##### Chimpanzee and gorilla

Whole brain samples from one chimpanzee (*Pan troglodytes*, 1 female, 54 years of age at time of death) and one gorilla (*Gorilla gorilla*, male, 12 years of age at time of death) were obtained from the London Zoological Society and the Primate Brain Bank. The samples were acquired from animals euthanized for management reasons unrelated to this research project. These brains were collected within minutes following death by initial perfusion with cold saline followed by cold paraformaldehyde. Details of sample preparation, scanning procedure, and data preprocessing have been communicated before ^36,37^. In brief, samples were stored in formalin, rehydrated using a phosphate-buffered saline solution one week prior to scanning and placed in fluorinert for the scanning procedure. Samples were imaged using a 7T whole body scanner with a 28-channel knee coil (QED). Diffusion MRI data were acquired using a 3D diffusion weighted steady state free precession (DW-SSFP) pulse sequence ^38^. DW-SSFP data comprising of 240 diffusion-weighted (q = 300 cm^-1^, resolution = 0.6 x 0.6 x 0.6 mm^3^) and six non-diffusion weighted (q = 20 cm^-1^) imaging volumes were acquired over the whole post mortem brains. To account for the T1, T2 and flip-angle dependencies of the DW-SSFP signal ^39,40^, T1, T2 and B1 datasets were acquired via a turbo inversion-recovery (TIR), turbo spin-echo (TSE), and actual flip angle imaging (AFI) acquisition ^41^. Prior to processing, a Gibbs ringing correction ^42^ was applied to the DW-SSFP, TIR and TSE datasets. Quantitative T1 and T2 maps were generated from the TIR and TSE datasets assuming mono-exponential signal evolution. A B1 map was generated from the AFI data following the methodology described in ^41^. All co-registrations within and between imaging modalities were performed with FLIRT ^43,44^ via a six degrees of freedom (translation and rotation) transformation. The DW-SSFP data, along with the T1, T2 and B1 maps were fitted with the full DW-SSFP signal equation ^39^ to both a diffusion tensor model and a ball-and-two-stick model using cuDIMOT ^45^.

##### Gibbon, macaque, squirrel monkey, and lemur data

Whole-brain samples were obtained from one gibbon (*H. lar*, male, 5.5 years old), three rhesus macaques (*M. mulatta*, between 11 and 15 years old, 1 female, 2 males), three squirrel monkeys (*S. boliviensis*, between 2 and 19 years old, 1 female, 2 males) and three ring-tailed lemurs (*L. catta*, between 3 and 11 years, 3 males). The samples were obtained from Copenhagen Zoo (gibbon, squirrel monkeys, and lemurs), the London Zoological Society (squirrel monkey), and the University of Oxford’s Biomedical Sciences (macaques). The samples were acquired from deceased animals that died of causes unrelated to this research project. Details of sample preparation, scanning procedure, and data preprocessing have been described previously ^46,47^. In brief, all brains were extracted and fixed within 24 h after the death of the animal. All brains were rehydrated in a PBS solution 1 week prior to scanning and placed in fomblin or fluorinert for the scanning procedure. The diffusion-weighted MRI data were acquired from the whole brain using a 7 T preclinical MRI scanner (Varian, Oxford UK). The scanner bore diameter is 210 mm, the gradient coil references are the following: 205_120_HD (Varian, Oxford UK) with a Gmax of 50 G/cm. The radiofrequency coil was made by Rapid Biomedical GmbH (Rimpar, Germany) and is a birdcage transmit receive coil with 72 mm ID. We used a 2D diffusion-weighted spin-echo multi-slice protocol with single line readout (DW-SEMS; TR = 10 s; TE = 26 ms; Matrix size = 128 × 128 with a sufficient number of slices to cover each brain; resolution for ring-tailed lemurs: 0.5 mm^3^; resolution for gibbon and macaques: 0.6 mm^3^; resolution for squirrel monkey 0.5 mm^3^. A total of 16 non-diffusion-weighted (b = 0 s/mm^2^) and 128 diffusion-weighted (b = 4000 s/mm^2^) volumes were acquired with diffusion encoding directions evenly distributed over the whole sphere (single shell protocol). Diffusion tensors were fitted to each voxel using DTIFIT ^48^. Subsequent preprocessing and analytical steps were specific for each species and are described in the following sections. Voxel-wise model fitting of diffusion orientations was performed using FSL’s BedpostX to fit a crossing fiber model to the data ^34^.

### Probabilistic tractography

Tractography was run using FSL’s probtrackx2 using the following parameters. Human: maximum of 3200 steps per sample; 10000 samples; step size of 0.25 mm; curvature threshold of 0.2. Macaque: maximum 3200 steps per sample; 10000 samples; step size of 0.1 mm; curvature threshold of 0.2. Chimpanzee and Gorilla: maximum 3200 steps per sample; 10000 samples; step size of 0.1 mm; curvature threshold of 0.2. Gibbon: maximum 3200 steps per sample; 10000 samples; step size of 0.1 mm; curvature threshold of 0.2. Squirrel Monkey: maximum 3200 steps per sample; 10000 samples; step size of 0.1 mm; curvature threshold of 0.2. Ring-tailed lemur: maximum 3200 steps per sample; 10000 samples; step size of 0.1 mm; curvature threshold of 0.2. It should be noted that diffusion MRI cannot distinguish between direct and indirect connections, so the identified fibers could represent either type of projection and travel in either direction.

### Comparative probabilistic tractography protocol

Probabilistic tractography of the anterior limb of the amygdalofugal pathway (AmF), uncinate fasciculus (UF), and the frontal segment of the cingulum bundle (CB) was performed in diffusion space. Descriptions of anatomical locations were made by reference to previously published atlases ^49,50^, tract-tracing studies ^21,51,52^, and diffusion tractography studies ^15,53^.

In all species seed, exclusion and waypoint masks were reconstructed in the same way with regard to established white and grey matter regions. The seed mask of the AmF was drawn in the sub-commissural white matter perforating the substantia innominata, a region between the dorsal amygdala and the bed nuclei of the stria terminalis (BNST). The mask was located medially to the ventral pallidum (VP) and sublenticular extended amygdala (SLEA) and dorsally to the nucleus basalis of Meynert (NBM) ^49,54^. Its position carefully aimed to reproduce macaque tract tracing studies ^51,55,56^. In accordance with the local trajectory of the AmF (as estimated by tract tracing), the seed mask was constrained to contiguous voxels showing high fractional anisotropy in an anterior-posterior direction. UF was seeded axially in the anterior temporal lobe, in the white matter rostro-lateral to the amygdala. An axial seed mask was chosen to account for the strong curve of the fibers from dorsal-ventral to anterior-posterior orientation to the orientation as they enter into the frontal lobe ^22^. The CB seed mask was drawn in a coronal plane capturing the WM dorsal and medial to the corpus callosum, within the cingulate gyrus. To increase the anatomical accuracy of our tracts, two additional tracts, the extreme/external capsule complex and the anterior limb of the internal capsule, were created and their seed masks used as exclusion regions for AmF, UF and CB. Excluding fibers related to these latter two tracts is important because they are expected to run or terminate in close proximity to AmF, UF and CB.

To constrain the tractography algorithm, exclusion masks were drawn in all species to exclude false-positive results in areas of high crossing fibers as follows: (1) within the basal ganglia to avoid picking up spurious subcortical tracts; (2) posterior to the seeds to prevent the projections from running backwards as the prefrontal cortical streamlines were the focus of this study; (3) an axial slice at the level of the superior temporal gyrus to prevent tracts from running in a ventral direction in an unconstrained manner (except for fibers reaching the amygdala); (4) on the axial-coronal slices cutting across the thalamus, basal ganglia, and corpus callosum to exclude subcortical and callosal projections; (5) in the dorsal cingulate cortex to avoid leakage of tracts to nearby bundles as a result of high curvature of the tracts (except for CB tracking); (6) an L-shaped coronal mask from the paracingulate cortex and the inferior frontal sulcus to the vertex to exclude tracking in the superior longitudinal fascicle; and (7) a mask excluding the opposite hemisphere to only track ipsilateral tracts. In humans, an exclusion mask of the CSF was used in each subject to prevent fibers from jumping across sulci during tracking. Streamlines encountering any of the exclusion masks were excluded from the tractography results. A coronal waypoint section was drawn in the frontal lobe at the level of the caudal genu of the corpus callosum to ensure that the fibers emanating from the seeds were projecting to the prefrontal cortex.

Streamlines were seeded from each voxel within each subject’s seed mask within the body of the tract of interest. We aimed to keep all seeding and tracking parameters constant across tracts and species, only adjusting step size based on brain size and white matter density. Each streamline followed local orientations sampled from the posterior distribution given by BedpostX ^34^. A visitation map or tractogram was constructed for each individual in order to allow comparison of these maps between tracts, subjects, and species. To counteract underweighting of distant voxels, the tractograms were log-transformed. Subsequently, to allow direct comparison across tracts, subjects, and species, independent of brain size and scan resolution, we further standardized the data by dividing a tract’s voxel values by the 75^th^ percentile value across the tract, thereby removing potential bias of differences in numbers of streamlines. In both human and macaques, the focus of the investigation was on the tracts’ connections with PFC in both hemispheres. If multiple subjects per species were available, the normalized tractogram was then warped to each species’ group template and averaged into a species-specific tractogram. For the purposes of visualization only, the normalized tractograms were subsequently thresholded. Figures were created with BioRender.com and FSL ^57^.

### Data sharing statement

Data were analyzed using tools from FSL ^57^, which is available online at fsl.fmrib.ox.ac.uk. Human MRI data were obtained from the Human Connectome Project (Van Essen et al., 2013) and available online from humanconnectome.org. The non-human primate MRI data are available from the Digital Brain Bank ^24^ online at open.win.ox.ac.uk/DigitalBrainBank.

## RESULTS

We reconstructed the principal connections between the limbic system and the prefrontal cortex (PFC) in the human brain and compared their organization with that in great (chimpanzee and gorilla) and lesser (gibbon) apes, an Old World (macaque monkey) and a New World monkey (squirrel monkey), and a prosimian (ring-tailed lemur). The evolutionary relationship between these species is shown in Figure 1. The goal was to assess how these connectional systems may have varied or remained consistent among primates. Our anatomical reconstruction focused on the temporal-prefrontal rostral branch of the amygdalofugal pathway (AmF), the uncinate fascicle (UF), and the anterior arm of the cingulum bundle (CB). We first describe reconstructions of all three tracts in the human and macaque monkey in detail, and then use them as baselines for subsequent descriptions of similarities and differences among the other species investigated. Importantly, here we defined anatomical and comparative approaches to reconstruct the anterior connections of the CB, AmF, and UF as they innervate frontal and temporal cortex. The approach used in each species were designed to be a similar as possible, allowing us to directly compare the organization of these fibre tracts within the methodological limitations. This follows prior studies that describe different brains in terms of directly comparable, homologous, features ^28,47,58,59^.

### Reconstruction of limbic-frontal connections in macaque and human

The UF is a C-shaped fiber bundle that reciprocally connects the temporopolar cortex and amygdala with the orbital and ventrolateral PFC. In our reconstruction, in a ventral section, the macaque UF extended along the outer surface of the temporal lobe and curved through the superior temporal gyrus (Figure 2). While the main direction of fibers in this region is from dorsal to ventral, some fibers branched along a medial-to-lateral axis towards the amygdala, thus connecting adjacent fibers near the anterior temporal region and the lateral prefrontal cortex (Figure 2). At the level of the limen insulae, UF fibers traveled in the white matter ventral to the claustrum, ventrolateral to the putamen and globus pallidus, and medial to the insular cortex (Figure 2). The human UF maintained a similar C-shaped structure projecting between the rostromedial temporal regions and the ventrolateral prefrontal cortex (Figure 2). Like the macaque temporal UF, it exhibited an anatomical organization primarily along a dorsal-ventral gradient, but with lateral fibers extending towards amygdaloid nuclei, parahippocampal areas, and rhinal areas (Figure 2).

**Figure 2.**
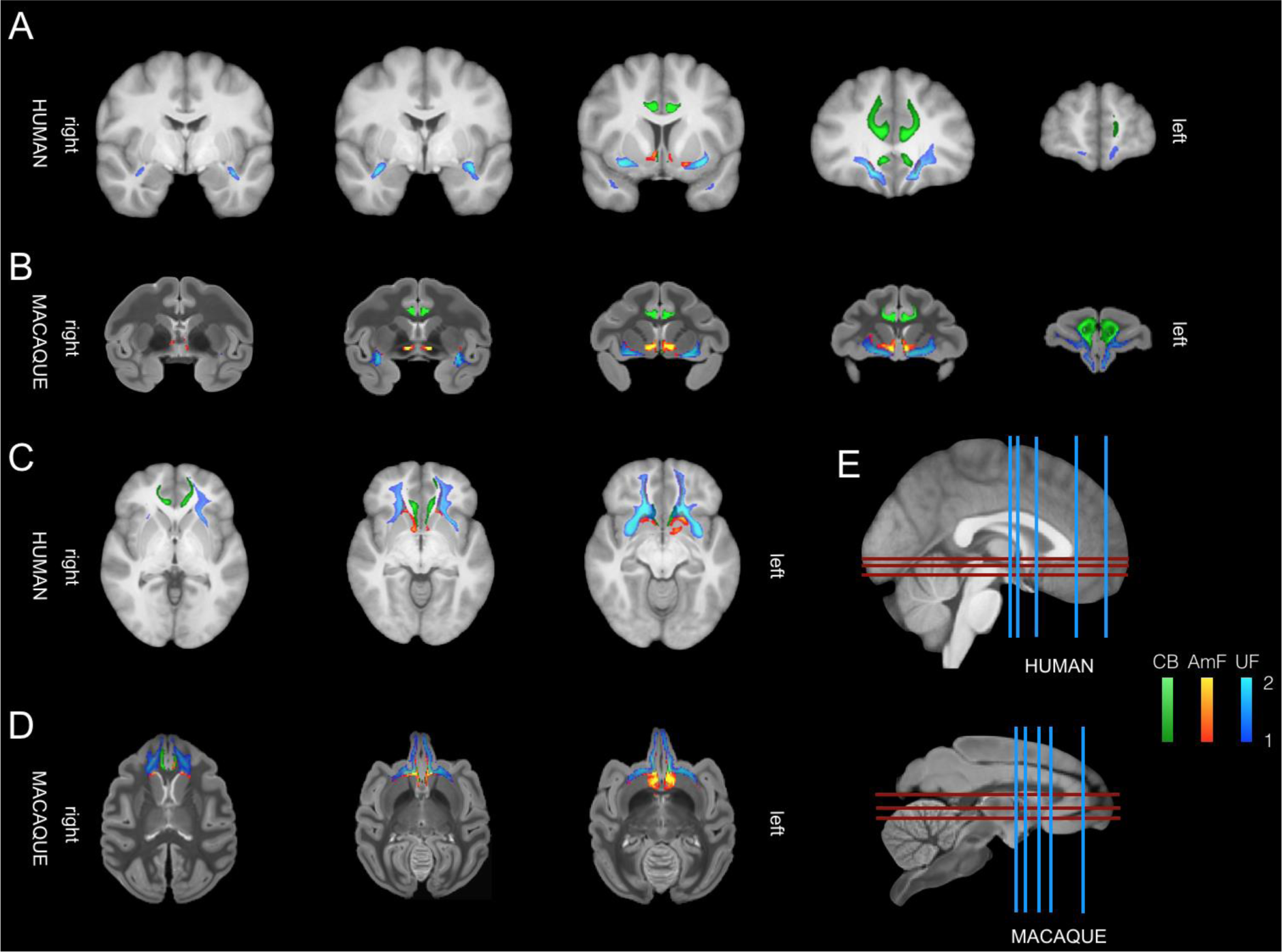
Limbic-frontal connections in macaque monkeys and humans. The anterior limb of the AmF (red-yellow), UF (blue-light blue) and CB (green) are displayed across the two species on five coronal brain slices (A-B) and three axial brain (C-D) slices showing their respective interactions and organization in limbic and prefrontal cortex. Location of slices from A-D are reported in E for the human brain (top) and macaque monkey brain (bottom).

Within the macaque prefrontal cortex, the UF exhibited strong connectivity with the white matter bordering the caudolateral orbitofrontal cortex (OFC), as well as opercular and inferior frontal areas. Additionally, a subset of fibers extended alongside the subgenual cingulate area and the medial bank of the posterior OFC (Figure 3-5). In this region, the more medial UF fibers joined with the fibers of the AmF and the CB to innervate the white matter of the anterior cingulate cortex (ACC) (Figure 3-5). Overall, the primary portion of the UF in the ventral prefrontal cortex spanned across the lateral, central, and medial orbital gyri, forming a complex network within the OFC that extended into the frontopolar cortex (Figure 2,4-5). Similarly, in the human, at the junction of the prefrontal and temporal cortices, the UF closely followed the insular cortex and areas within the caudal region of the inferior frontal gyrus (Figure 2). Along the basal ganglia, the human UF encompasses the lateral ventral striatum and extends to a more lateral section of the ventral prefrontal cortex (Figure 2). In this region, the main body of the macaque and human UF continues to traverse laterally, providing extensive innervation to the white matter of the ventrolateral prefrontal cortex. Further, a subset of fibers also courses more medially, adjacent to subgenual anterior cingulate cortex (sACC) and, more prominently, towards the posterior OFC alongside the AmF and the CB (Figure 3-5). Within the anterior region of the ventral prefrontal cortex, both macaque and human UF extends across medial and lateral orbital areas. A significant bundle of fibers occupies a substantial portion of the white matter in the lateral OFC and adjacent cortical areas on the ventral surface of the brain in both species. Another branch of the UF extends further rostrally in the white matter adjacent to the frontopolar cortex (FPC) in both macaque and human brains (Figure 2,4-5).

**Figure 3.**
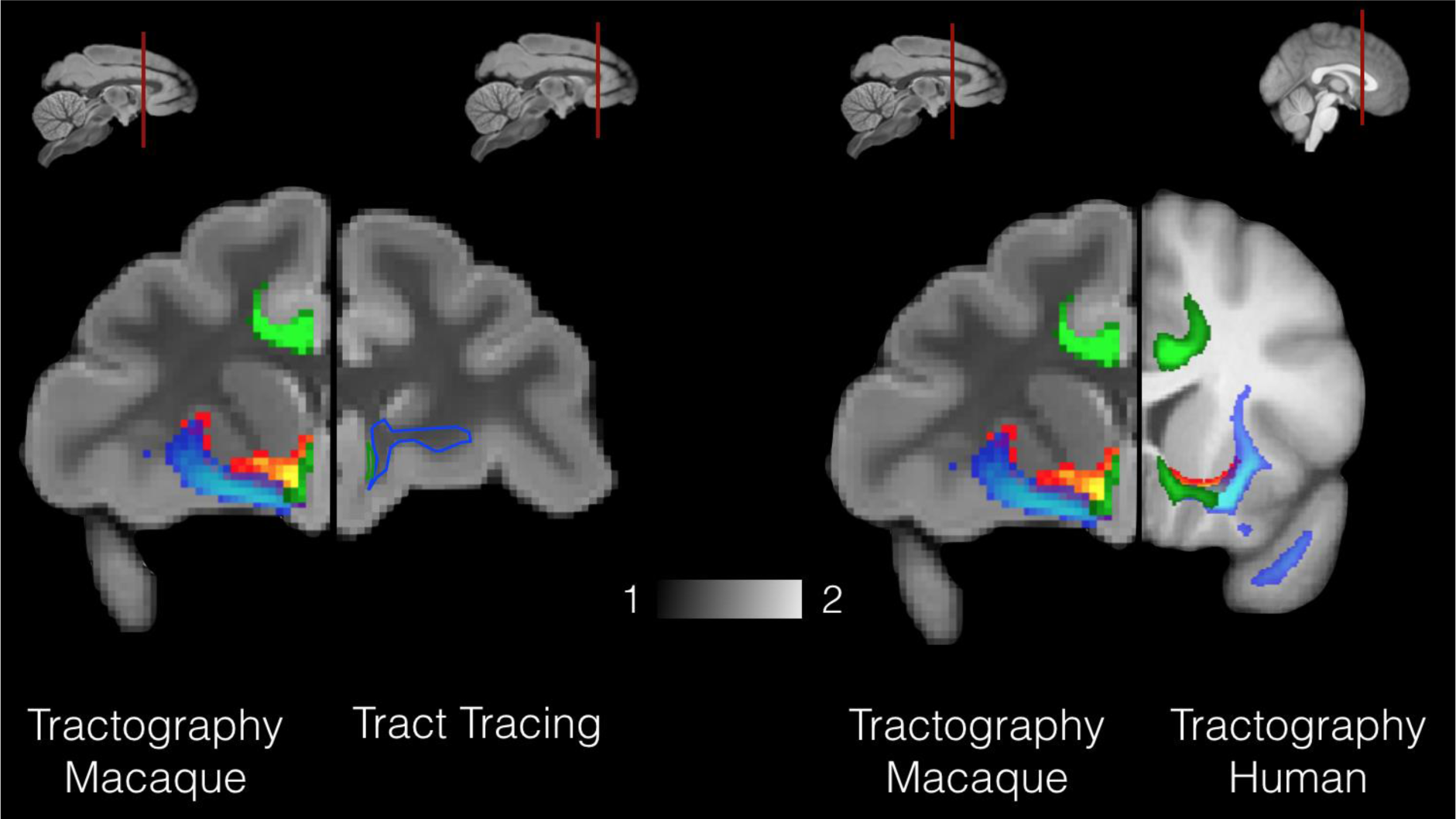
Histology-diffusion MRI comparison of connections. Tract-tracing-driven (left panel; reconstructed from Figure 15 in Heilbronner and Haber^63^ reconstruction of amygdala-cingulate-prefrontal connections using non-invasive diffusion MRI (left and right) in macaque monkeys (right hemisphere) and humans (left hemisphere). In both macaque and human brains AmF (red-yellow), UF (blue-light blue) and CB (green) are shown.

**Figure 4.**
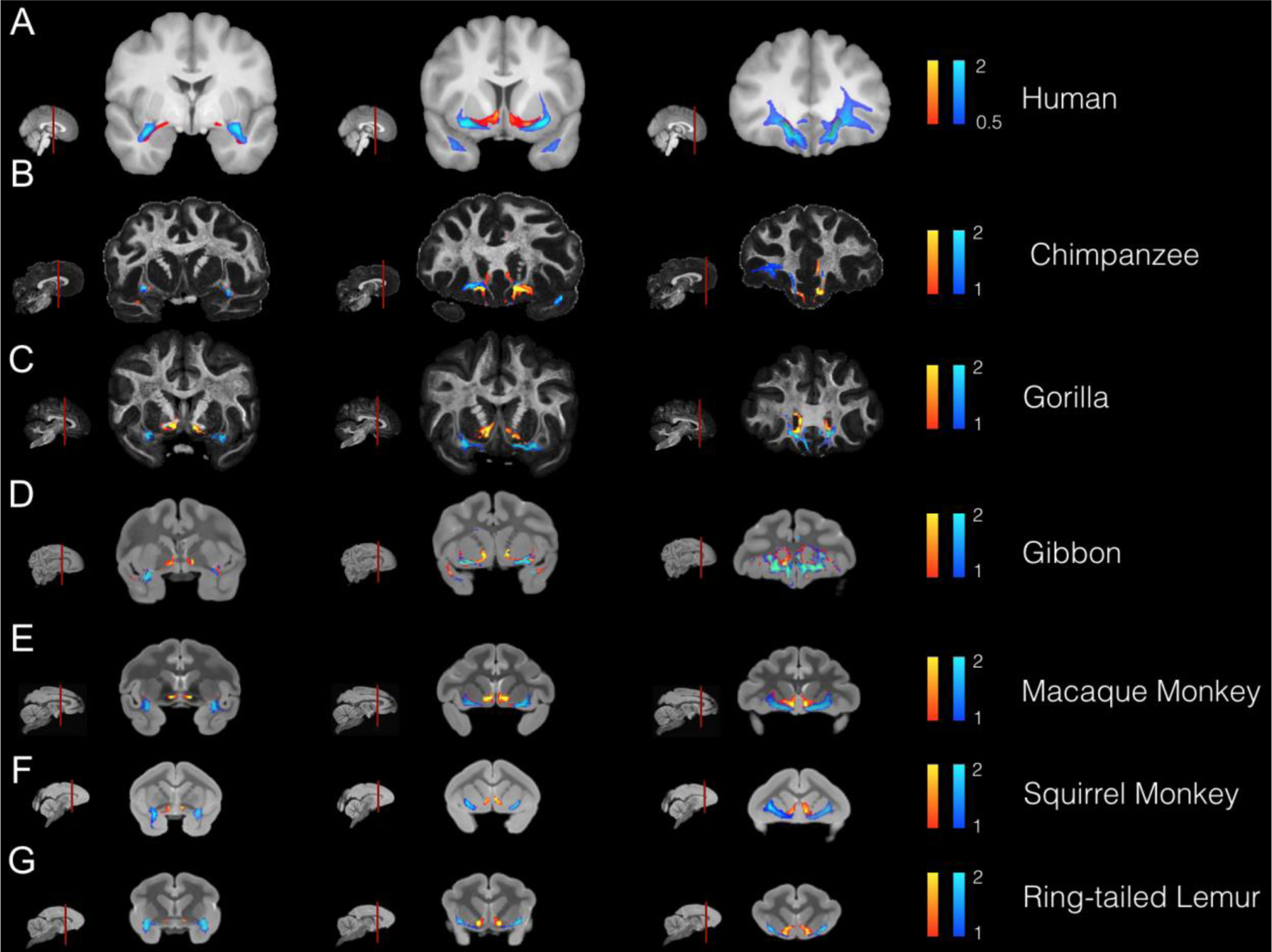
Coronal views of temporal-frontal connections in all species. The anterior limb of the AmF (red-yellow), UF (blue-light blue) are displayed across each species on three coronal brain slices showing their respective interactions and organization along a posterior (leftmost column) – anterior (rightmost column) gradient. The brains are displayed from the species most closely related to humans (top row) to the least related (Ring-tailed lemur, bottom row).

**Figure 5.**
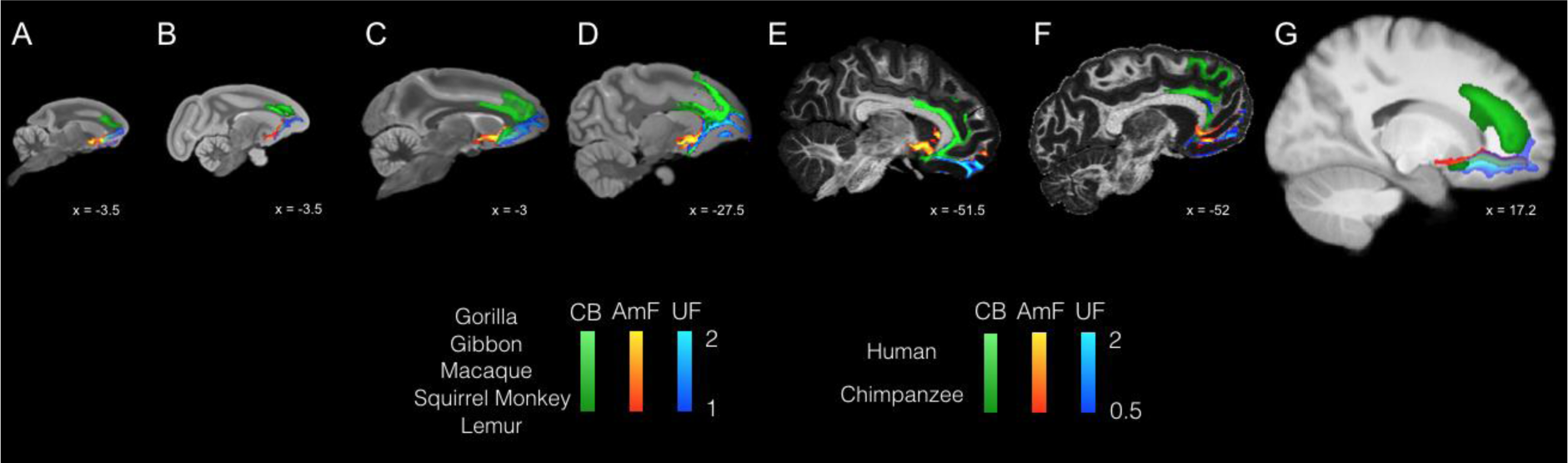
Organization of temporal-cingulate-frontal connections across all species. The anterior limb of the AmF (red-yellow), UF (blue-light blue) and CB (green) are displayed across each species on a sagittal brain slice showing their organization within cingulate and prefrontal cortices, striatum and basal forebrain. The brains are displayed in the following order from left to right: ring-tailed lemur (prosimians), squirrel monkey (New World monkey), macaque monkey (Old World monkey), gibbon (lesser ape), gorilla (great ape), chimpanzee (great ape) and human (*Homo sapiens sapiens*).

In the macaque brain, the frontal portion of the AmF was represented by a thin group of fibers that ran in the white matter adjacent the longitudinal fissure. At the level of the anterior commissure (AC), the fibers in the medially projecting limb of the AmF pathway traveled within the white matter dorsal to the amygdala, between amygdala nuclei, sublenticular extended amygdala area and medial longitudinal fissure (Figure 2,4). In this species, one set of AmF fibers ran in the white matter region between the nucleus basalis of Meynert, the lentiform nucleus, the ventral pallidum, and the piriform cortex. These medial AmF fibers divided into different sets of projections. Some extended downward to innervate the anterior hypothalamic areas near the paraventricular-supraoptic nuclei, while others coursed dorsally to connect the amygdala with the thalamus, the BNST, and the septum of the macaque monkey. Another set of macaque AmF fibers ran in the white matter surrounding the substantia innominata, ventral pallidum, and the lower bank of the AC, entering, further rostrally, into the white matter beneath the ventral striatum/nucleus accumbens (Figure 2,4). These AmF pathways were found in both macaque monkeys and humans. Despite the strong medial-lateral primary diffusion direction characterizing the AC, some weaker AmF fibers that overlapped with the AC were still detectable in our tractography data within both macaque monkey and human brains. It is important to note that as the AmF passes through the basal forebrain and beneath the striatum, it is likely to interact with other connectional systems.

Eventually, the anterior fibers of the AmF of macaques and humans extended into the orbital cortex and into the rostral PFC by running in the white matter adjacent to the subgenual anterior cingulate cortex (sACC or Brodmann area 25) and the adjacent areas in subcallosal cortex (Figure 3-5). Near the sACC, the AmF of macaques and humans split into two subsections. The first subsection ran upward alongside the CB in the anterior cingulate sulcus and the surrounding white matter (Figure 5). The second branch, on the other hand, stretched between the sACC and the frontopolar cortex, passing underneath the medial OFC between the gyrus rectus and the medial orbital gyrus (Figure 2,4-5).

Frontal connections of the macaque and human CB project along the full extent of the anterior cingulate cortex. This bundle is composed by a mixture of fibers entering the large cingulum bundle briefly to reach nearby areas or longer-distance connections reaching distal brain regions. Core CB fibers surround the corpus callosum throughout its course in PFC in both species. A subsection of fibers in rostral CB exits the main bundle and contains axons towards and arriving from both cortical and subcortical areas (Figure 5). At the level of subcallosal and orbitofrontal cortex, in both macaques and humans, CB fibers interact with axons travelling in the AmF and UF (Figure 3,5). Here, CB fibers either terminate by innervating adjacent areas or continue coursing in a rostral direction towards anterior PFC and frontopolar cortex. In both species, another set of connections continue running along the genu of the corpus callosum, next to area 23, 24 and 32, and more dorsally, in the CB branches extending into dorsal ACC regions, premotor, lateral PFC (partly through intermixing with UF fibers) and medial-dorsal PFC areas 8 and 9 (Figure 5).

Overall, these results, obtained by probabilistic tractography accounting for multiple possible fiber orientations ^34^, show that AmF, UF and prefrontal CB fibers exhibit a high degree of similarity between macaque monkeys and humans (Figure 3-4). In both species, fibers from the three bundles merge and intertwine at the level of the subgenual cingulate cortex giving before partially separating and running independently in anterior prefrontal and temporal cortices (Figure 3-5). The protocols employed here to reconstruct UF, AmF, and CB in macaque and human were based on Folloni et al. ^22^, tailoring them to obtain results best matching those previously observed using macaque tracer data ^63,64^. This enabled us to avoid false positives and false negatives that may be created when employing tractography algorithms and ensured the truthfulness of the reconstructed bundles. Figure 3 shows the macaque and human tractography results side by side with previously published tracer data to corroborate these original findings. Having established a protocol that allowed both matching of macaque tractography data with established macaque tracer data and matching of macaque and human data, we felt confident using these protocols in chimpanzee, gorilla, gibbon, squirrel monkey, and ring-tailed lemur. We describe the course of the tracts in these species at the levels of anterior temporal cortex, subcallosal frontal cortex, and more anterior prefrontal cortex.

### Course of the AmF and UF between temporal and frontal cortex across all species

The most prominent shared feature observed across lemur, squirrel and macaque monkey, gibbon, gorilla, chimpanzee and human brains is their widespread connectivity between prefrontal and anterior temporal areas, both subcortical medial and lateral cortical structures (Figure 4). Consistently across all the species examined, fibers connect amygdala, piriform and inferior temporal cortices on the temporal side with medial and lateral regions in anterior PFC on the opposite end. As the anterior limb of the AmF and the UF travel between temporal and frontal poles, these bundles innervate striatal, insular, basal forebrain, ventromedial and ventrolateral PFC areas (Figure 2). Consistently across all the species examined, the AmF exits the amygdala via the medial wall of the temporal lobe and then runs dorsally towards limbic, striatal and prefrontal structures along other forebrain and thalamic fibers (Figure 2. The UF, instead, unfolds as a curve between the anterior temporal lobe and PFC carrying prefrontal, forebrain, amygdalar, rhinal, piriform, parahippocampal and temporal associative projections.

A clear distinction between AmF and UF is observable across the six primate species at the level of the amygdala, temporal cortex, and peri-commissural white matter, showing the dichotomous medial-lateral organization in striatal and basal forebrain we see in macaque and human (Figure 4). The AmF occupies the white matter perforating the ventral pallidum and expanding underneath the ventral striatum, primarily branching towards the anterior ventromedial striatum, rostrally, and the hypothalamus, bed nuclei, stria terminalis and septum, medially. UF fibers course through the sub-capsular white matter along the ventral insular cortex and putamen. A few fibers leave the main body of the UF and merge with the AmF, approaching from a central position. This ventral-dorsal intertwining of the two paths is observable across all primate brains examined and becomes complete when AmF and UF approach the subgenual ACC (Figure 2). In the white matter innervating the striatal and basal forebrain territory, however, AmF and UF in the gibbon overlapped to a higher degree than in the other species, where they were still clearly distinguishable (Figure 2). Interestingly, the medial-lateral organization of UF and AmF is also preserved in the strepsirrhine lemur (Figure 2) despite the anatomical expansion of the temporal lobe and of the PFC in anthropoids^60,61,65,66^.

### Level of the subcallosal cortex

Fibers coursing alongside the subgenual ACC, underneath the ventral floor of the genus of the corpus callosum, display a common organization across Old World monkey, lesser and great ape, and human brains. After coursing through the temporal lobe and near the ventral striatum, AmF and UF approach the subgenual cortex from a medial and lateral position respectively. This is also where the main body of the cingulum bundle (CB) joins the territory of AmF and UF, but its main body runs primarily medial to AmF and UF (Figure 3,5). In both the macaque and human brains, the CB fibers merge with fibers from AF and UF in this region, confirming results from previous tract-tracing studies ^67^.

In caudal subgenual ACC, AmF and UF not only begin sharing important portions of white matter with CB, but they also start merging into a single limbic bundle innervating ventral PFC and anterior temporal cortex, in addition to medial cortex at their extremes. Although AmF and UF here begin merging into a single connectional system as they interacted with CB they maintained a medio-lateral organisation until they approach posterior OFC. The most dorso-medial WM territory surrounding the subgenual ACC is primarily occupied by fibers carried by an additional dorsal limbic tract, the CB. Within the PFC, this tract carries fibers running in the middle, anterior and subgenual cingulate cortex.

Interestingly, CB fibers did not extend into the caudal sACC as clearly as in the Prosimian (ring-tailed lemur) and New World monkey (squirrel monkey) but we could still observe a similar organization of AmF and UF as described above for the macaque monkey, gibbon, gorilla, chimpanzee and human brains. This pattern was consistent even when adjusting for different values of the curvature parameter used in our analyses aiming to account for the anatomy of the genu of the corpus callosum in these two species. (Figure 2).

### Level of anterior prefrontal cortex

Thus far, the organization of UF, AmF, and CB is similar across all of the primates studied here. However, prefrontal cortex has differentially expanded in different primate lineages, showing a relative increase in size in anthropoid primates (Passingham and Wise, 2012) and perhaps again in the ape and especially the human lineage ^26,27^. It is therefore of particular interest to examine how these tracts project to anterior prefrontal cortex.

A merging of amygdala, temporal cingulate and prefrontal connections in posterior OFC was clearly observable along all of the representatives of the primate order examined (Figure 4). As previously reported for macaque and human brains ^22^, AmF and UF transition from a dichotomous organization in temporal and subcortical regions to an intertwined bundle in more anterior PFC white matter. Although in medial portions of anterior PFC, AmF, and UF occupy a similar portion of white matter, we observed that AmF occupies the territory adjacent to the medial frontal wall, whereas UF extends to lateral ventral areas.

In anterior PFC, AmF, and UF fibers coursed together along the medial orbital and frontopolar cortices consistently across lemurs, squirrel and macaque monkeys, gibbons, gorilla, chimpanzees and humans regardless of their frontal lobe expansion (Figure 4). In addition, UF fibers branched out to caudolateral OFC, ventro-lateral PFC as well as opercular and inferior frontal areas (Figure 4). Axons from both AmF and UF reached granular PFC in the primate species that possess this cell type. Anatomical interactions among AmF, UF and CB fibers in caudal-ventral PFC were shared across all species, except in the gibbon, where AmF and UF converged more posteriorly, beginning in the transition territory between subgenual and orbital cortex. Although we cannot entirely rule out the influence of false positives given the small sample size for this species, it is noticeable that the gibbon is the only species among those examined here that primarily lives in very small groups, and spends the least amount of time in socially and emotionally bonding behaviors ^68,69^. This peculiarity in social behavior, either it being environmentally induced or inherent in the gibbon nature, may be associated with our finding that there is a difference in AmF and UF organization in posterior OFC for this species.

## DISCUSSION

In this study, we reconstructed the limbic-prefrontal connections in species from all major branches of the primate radiation. By using the same methodology, we showed that the overall organization of AmF, UF, and CB is evolutionarily conserved, but with some species-specific variances. This is an important result in light of the current need for improved translation of findings in animal models to humans to facilitate the development of novel brain treatments. Importantly, our results from diffusion tractography reveal that the anatomy of limbic and prefrontal fibers resembled previous anatomical tracer investigations focused on these specific tracts in macaque monkeys ^21,52,56,63,64,70^ and diffusion imaging studies that compared only a subset of the species investigated herein ^22,36,71^.

Primate brains, including those of humans, all follow a general organizational plan, but each brain is expected to show individual adaptations ^72^. Both temporal and frontal cortex have undergone expansions in the lineage leading to apes and humans ^27,73,74^, accompanied by an increase in organizational complexity and differentiation of both the grey and the white matter ^17,36,75^. This is, to our knowledge, the first study that compares the organization of a set of specific white matter tracts across all major branches of the primate family tree. Earlier work has either focuses on more global measures of connectivity ^76,77^ or looked at a limited set of species ^17,36^.

Despite the aforementioned anatomical heterogeneities in temporal and frontal cortex across primates, it is intriguing that the overall organization of AmF, UF, and CB appears to be conserved. Specifically, across all species, amygdala, anterior temporal cortex and prefrontal cortex connections travel in three separate fiber tracts--the AmF, the UF, and the CB—that show a similar course and similar topology with respect to one another. This suggests that the organization of the AmF, UF, and CB represents a shared anatomical feature, possibly originating in early primates. Inter-species similarities in connectivity within the anterior temporal and frontal lobes may be indicative of how the environment shaped white matter organization in early primates. Further, these shared organizational features may be implicated in the development of shared foraging strategies across this order, that primarily rely on visual foraging and associative learning of visual stimuli ^6,13,66,78–85^.

It is important to properly contextualize this result. We show that the organization of AmF and UF fibers in the basal forebrain and subgenual cortex in dissociable pathways carrying limbic-prefrontal fibers is conserved across primates. This is an especially intriguing finding in lemurs and squirrel monkeys, due to the remarkably smaller density in white matter within this region in these two species. Our study methodology focused on reconstructing the main bodies of known fiber bundles throughout the brain. This has been shown to be a particularly reliable way of comparing aspects of white matter architecture across species, as it allows identification of anatomically similar structures across species ^28^ and, unlike inferences regarding the connectivity of specific grey matter areas, is not susceptible to false positives ^86^. This approach is therefore the basis of tools for comparing white matter architecture across species, development, and individuals ^87,88^. However, this does mean we do make claims on the precise grey matter areas innervated by these bundles. Indeed, the conserved fiber systems identified in this work exist within neural systems that do show phylogenetic diversity, as mentioned above. A particularly prominent difference is the addition of granular areas in the prefrontal cortex ^58^. Ongoing work will determine how these converged pathways relate to diverse grey matter architecture ^60^ and the implications of this for function ^89^.

Despite the overall conservation, our results did suggest a potential differences in innervation of subgenual cortex across species. Although still poorly understood, subgenual cortex is emerging as a key hub for mood and affective processing, as dysregulation of this region is a core feature of patients with mood disorders and anhedonia ^90,91^. Lesions of this area cause deficits in anticipatory arousal ^92^, whereas hyperactivation is associated with reward processing deficits and autonomic changes ^93^ as well as impairments in aversive behavior ^93,94^. These findings implicate subgenual cortex as critical to the orchestration of neural circuits regulating affective behavior and survival ^95^. In most species, the cingulum bundle interacts in subgenual cortex with AmF and UF before the three bundles separate into their own specific trajectories. These results replicate tracer studies in Old World monkeys ^63,78,79^. We did not observe the exact same organization in the lemur and squirrel monkey, as none of the reconstructed cingulum bundle fibers, when thresholded at the same level as the other species, were able to curl around the genu of the corpus callosum and reach the subcallosal white matter, regardless of the curvature parameter we used.

It is important to consider what such a result means. It is unlikely that no part of the cingulum bundle projects to this caudal part of subgenual and ventromedial PFC in these species, as also shown by a tracer study in the squirrel monkey that demonstrated the existence of cingulate fibers coursing in this portion of WM ^96^. However, the density of fibers reaching this territory, as indicated indirectly by the likelihood of the probabilistic tractography reaching there, does seem to be much lower in these species. This is important, as the density of connections is a very important predictor of information exchange between areas ^97^. Our results suggest that the increasing use of New World monkeys such as the squirrel monkey in translational research ^98^ should be accompanied by extensive validation studies to ensure that results obtained in other model species replicate.

In conclusion, we show that despite the variations in phylogenetic distance, life history, behavior, and the environments inhabited by the species in our study, the topology of anatomical connections between the limbic system and frontal cortex is shared not only across humans, apes, and monkeys, but also with simians and prosimians. Although these limbic-frontal connections innervate systems that show variation in size and morphology, their overall architecture and course is conserved. An exception to this are the cingulum connections to subgenual cortex, which is an important target for neuropsychiatric treatment. Together, this work shows how a large-scale comparative approach can help guide translational neuroscience ^99^, as well as provide insights into brain evolution.

## Acknowledgements

The work of RBM was funded by the Biotechnology and Biological Sciences Research Council (BBSRC) UK [BB/N019814/1]. The Wellcome Centre for Integrative Neuroimaging is supported by core funding from the Wellcome Trust [203139/Z/16/Z]. The work of KLB was funded by Institut Convergence ILCB (ANR-16-CONV-0002). For the purpose of Open Access, the authors have applied a CC BY NC ND public copyright license to any Author Accepted Manuscript version arising from this submission.

## Author contributions

D.F., R.B.M. planned and designed the study. D.F., R.B.M., P.M., M.F.B., A.A.K., acquired the data. D.F., R.B.M., L.R., K.L.B. analyzed the data. D.F., R.B.M., K.L.B., P.H.R., interpreted the results. All authors contributed to manuscript preparation.

